# How Many Females Are There? Cheating and Female Dispersion Can Explain Mating Behavior Evolution

**DOI:** 10.1101/171330

**Authors:** B. V. Gomes, D. M. Guimarães, D. Szczupak, K. Neves

## Abstract

Only around 3% of all mammalian species are socially monogamous and the conditions that favor the evolution of this mating system in mammals are not well understood. With several approaches, studies have proposed different hypotheses relating female dispersion and infanticide as drivers for the evolution of social monogamy. Here, we used an agent-based model, that allowed us to examine how different mating behaviors affect populations in a controlled computational environment. We found that the evolution of social monogamy does not rely on a single factor. Rather, our experimental results support an interplay of different factors in the evolution of social monogamy – female dispersion and availability and breeding season duration – and suggests that polygamy will only evolve in populations with a female-biased operational sex ratio or one where cheating is common. These results can explain why social monogamy is so uncommon in mammals and points to new lines for ethological investigation of mammalian behavior.

---

The most recent consensus defines social monogamy as the sexual engagement of a male and a female for more than one breeding season (*1*). This form of mating system is common in birds, but very rare in mammals (*2*). Social monogamy is intriguing because monogamous males invest time in its exclusive partner, time that could be used to mate with several females, which should increase its fitness. Which selective pressures lead to monogamous behavior in mammals? Why is polygamy the norm in mammals? Under which conditions should we expect monogamous or polygamous behavior to be advantageous?

In the last decades, researchers have proposed many hypotheses to explain how social monogamy evolved in particular cases, for example, in mammals (*3*, *4*). Of all of them, two noteworthy hypotheses relate to male infanticide (*4*, *5*) and female dispersion (*6*, *7*). To understand the intricate play of factors, scientific efforts have used two experimental approaches: either generalizing based on scarce ethological observations (*8*-*10*) or reconstructing phylogenetic trees based on ancestral behaviors and inferring the causes of evolution of social monogamy based on order of appearance (*11*, *12*). Both approaches rely on the observation and categorization of complex behaviors, a dauntingly hard task. Because of a lack of consensus about how these steps are to be done, evidence is contradictory (for an example of two contrasting views, see *5*, *6*, *13*, *14*).

Others have used equation-based models (EBMs) to model large-scale mechanisms and study the fitness of different mating strategies (*15*-*17*). Usually, they consider the relationships between the variables, and abstract the specific empirical data, except to ground the model in reality. They are useful for understanding mechanisms and clarifying assumptions, but are met with skepticism by “ethological purists„ (*18*). In any case, they have clear limitations: in particular, EBMs are more suited to study systems where local spatial dynamics are not relevant and where individual-level behavior does not vary much. In systems where there is local variation and heterogeneity of individual behavior, agent-based models (ABMs) are more appropriate (*19*). Agent-based modeling is a strategy that focuses on modeling individuals and their interactions instead of system-level observables. It defines the individual agents and simulates how it evolves over time, letting emergent or system-level properties to appear from the bottom-up. It allows a researcher to set specific conditions and study how particular behaviors could differently affect outcomes, allowing experimentation and more direct translation to the actual phenomena, when compared to EBM. In recent years, this modeling approach has brought new insights to the study of ecology in general (*20*) and even the evolution of monogamy (*21*).

Taking into consideration the hypotheses put forward in the literature to explain the evolution of mating behavior in mammals, we wanted to see if changes in the behavior of individuals, induced by local changes in their circumstances -external attributes, both social and ecological (*22*) -can lead to the evolution of a polygamous or monogamous phenotype in the population. In particular, to investigate the female dispersion hypothesis and better understand the underlying mechanisms and circumstances where the hypothesis holds, we developed an agent-based model of the evolution of a generic species and its breeding system. In our ABM, males move around searching potential mates. They are either monogamous or polygamous – which determines how they search for females – and this tendency is treated as a genetic trait which is passed to the next generation. We then follow the spread of monogamous or polygamous behaviors in the population of male agents as a primary outcome, while changing parameters of interest across simulations to test our hypotheses.

## Results

### Female dispersion hypothesis

Our results show that female dispersion does have an effect, in the sense predicted by the hypothesis (Figure 1) – the larger the female dispersion, the more monogamous the population. However, this effect is only present in some circumstances. If the number of males is equal to or larger than that of females, there is a strong tendency to monogamy, irrespective of the dispersion. That not being the case, then if the duration of the breeding season is long, there is a strong tendency for polygamy. When neither is true, female dispersion is a good predictor of monogamous behavior, like expected by the hypothesis (Figure 1, red data points, R^2^ = 0.75, for simulations with few male agents (15) and a short season duration (200 steps)).

**Figure 1.**
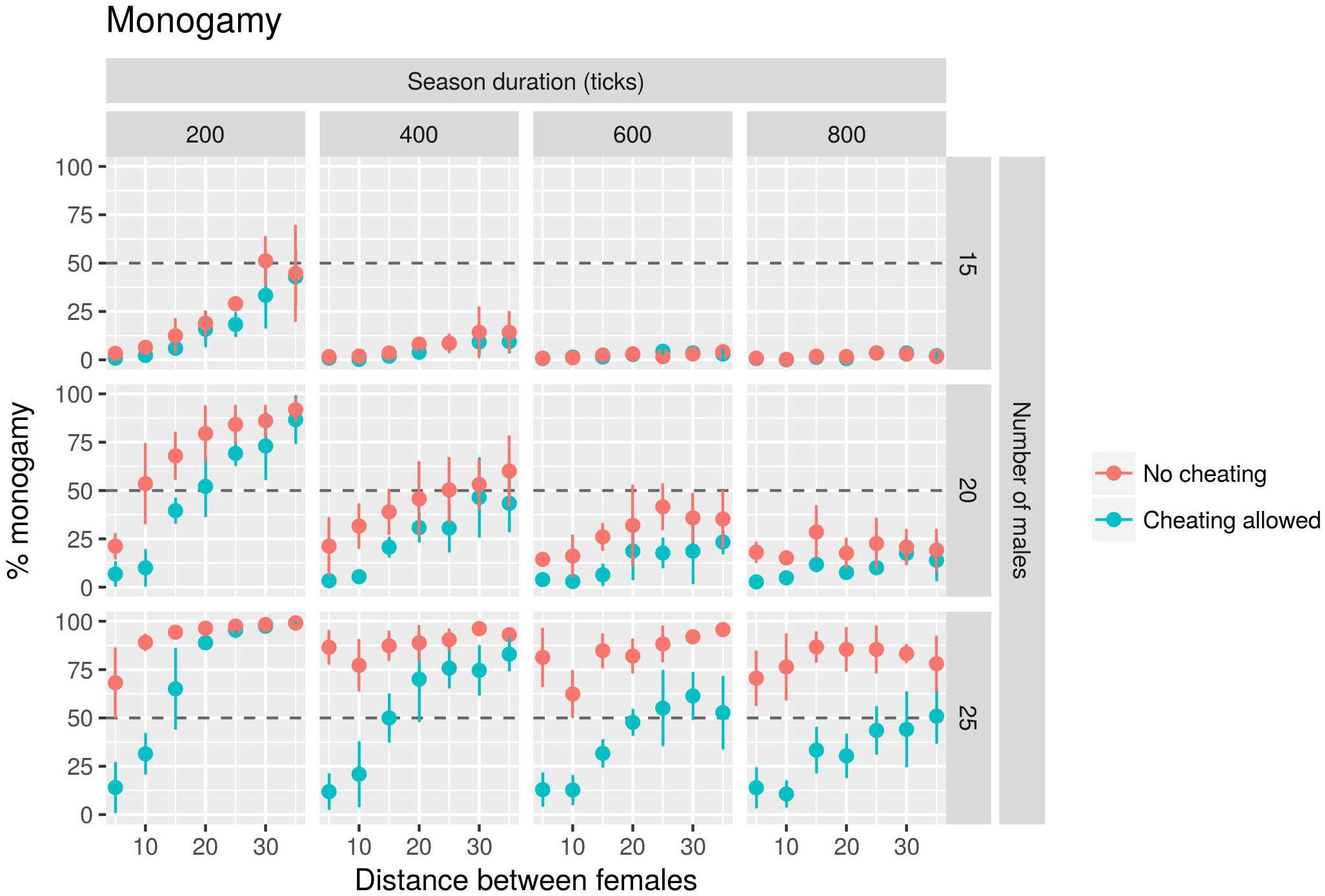
Monogamy dynamics varies according to the interaction of parameters. Each point shows the mean percentage of monogamy (the percentage of seasons where over half the population of males was monogamous, see Methods Section; n = 5 replicates) and bars show 95% confidence intervals. The traced line indicates 50% of monogamy. Simulations were performed with 20 females (arbitrarily chosen), distance between females varied from 5 and 35 patches (**bottom axis**), breeding season varied from 200 to 800 ticks (**columns**), number of males varied from 15 to 25 (**rows**). Percentage monogamy (**left axis**) positively correlates with female distance (**top left graph**) and number of males (**note last column**) and negatively correlates with season duration (**note the first row**) and the presence of cheating (**notable in every graph**).

We interpret these results as follows: initially, monogamous and polygamous males are equal, all of them searching for females. However, once a monogamous male finds a female, pregnancy is almost guaranteed – given a long enough season. Polygamous males, on the other hand, continue copulating after the point where most monogamous males have stopped – because their female partners are pregnant. The longer the breeding season, the longer the polygamous males will continue copulating beyond the monogamous ones. For a shorter season, there is no time for polygamous males to compensate an initial advantage of monogamous males in case females are very dispersed, therefore, the distance between females has a stronger effect on the fitness for each behavior. Moreover, this compensation can only happen if there are enough females available, which explains why polygamy only appears consistently in situations where females outnumber males.

### Operational sex ratios and extrapair copulation

Based on the previous results, if polygamy were to be a common mating system, we would expect that the operational sex ratio (OSR) to be female-biased in most mammals. The OSR -defined as the ratio of sexually active males, ready to mate to females that are sexually active, ready to mate -is the ecological measure most closely related to the number of male and female agents as represented in our model. While, to our knowledge, there is no systematic analysis of OSR in mammals, it is known from the literature shows that female-biased OSRs are not the rule (*23*, *24*), therefore it doesn’t hold as a general explanation.

Another possibility is that males that cannot find females in their current location can generally move to another location where available females can be found. However, there was yet another, more interesting, possibility: that females paired with monogamous males are not necessarily unavailable for polygamous males, because extrapair copulation may occur (*10, 25*). This would effectively increase the number of available females at any given moment and remove the guarantees monogamous males had of a pregnancy once they find a suitable female.

To test this, we made a simple change to the ABM: monogamous males no longer stayed beside their females at all times, allowing the possibility of extrapair copulation (see Methods for details of implementation).

Allowing cheating had the expected effect: when we decrease the monogamous male protection and allow cheating, we see the effect of breeding season duration and female dispersion on mating behavior even when males outnumber females. The dynamics become even clearer, fitting our interpretation: monogamy is a better strategy when (1) there are few females available relative to males, (2) when the reproductive season duration is short and (3) when female dispersion is high.

### Regression model results

Finally, to confirm our intuitions, we ran a regression of our primary outcome – the percentage of time when over half the population was monogamous – on number of males, breeding season duration, female dispersion and a dummy variable indicating whether or not cheating was present. The results show a good fit for the full model (adjusted R^2^ = 0.78, p < 2. 10^-16^) better than models fit with each variable in isolation (female dispersion: R^2^ = 0.21, p < 2. 10^-16^; season duration: R^2^ = 0.15, p < 2. 10^-16^; number of males: R^2^ = 0.36, p < 2. 10^-16^). Interestingly, the regression shows that the predictors with larger effects are the proportion of males (0.602) and female dispersion (0.310), followed by season duration and cheating presence, which have a negative effect on monogamy (-0.288 and -0.185, respectively, all p-values < 2. 10^-16^).

## Discussion

We built an agent-based model to study the spatial dynamics of monogamous and polygamous behavior. Using a computational model, we were able to experiment with different ecological parameters – parameters which prior hypothesis and empirical results suggest might have an effect on the evolution of mating behavior – to see how they affect the spread of different mating strategies in the population. This sets up a platform to study of the interaction of different factors in the evolution of mating behavior, which is otherwise difficult to do, be it in verbal models or in empirical studies, due to the sheer amount of elements and forces involved in the system or the impracticality of manipulating large populations experimentally. The model could be easily extended in future work to investigate the effect of aspects that we didn’t explore in this work but which are featured in the literature, such as infanticide (*5*) or parental care (*6*).

We show that polygamy is only possible when either (1) the OSR is female-biased or (2) extrapair copulation is widespread among females or both. Since polygamy is the norm in mammals (6), one suggestion that comes out of our model is that most mammals have female-biased OSRs. While available evidence does not seem to support the generality of this affirmation for all mammals (for instance, see 23, 24), it remains to be systematically investigated. Nevertheless, studies with a crustacean, the snapping shrimp (*Alpheus angulatus*) showed that in conditions of female-biased sex ratios, males were more likely to search for new females after mating (*26*) The other suggested possibility is that cheating behavior is common in socially monogamous mammals. This proposition finds some support in recent studies showing that animals assumed to be genetically monogamous are only socially monogamous – this is the case of the prairie vole (*Microtus ochrogaster*), once held as a model of lifelong sexual fidelity (10). It’s also known that prairie voles which engage in extra-pair copulation have a larger range and venture outside their home territories more often (*25*) Our results reinforce the case for a separation between social bonds and sexual fidelity in mammals. Moreover, they point to sexual fidelity as a driver of higher female availability, which in turn favors polygamous strategies, according to the dynamics in our model.

In accordance with recent literature (6), we conclude that female dispersion is a viable mechanism to explain the evolution of monogamy in mammals, albeit only in specific conditions. Our model revealed other important factors in the evolution of mating behavior. First, availability of females is important, but not only conceived as local availability influenced by dispersion, also as the sex ratio, which has consequences for the competition between males and influences the cost of roaming – here represented minimally as time lost, but which could also involve risk of aggression by other males, for instance. One other parameter, not directly related to female availability, also has an effect: the duration of the breeding season. We reason that in a short breeding season monogamous behavior is advantageous because the shorter the season, the less likely it is that a male will find more than one female, which means leaving the one female unguarded has no advantage. While the effect of breeding season duration was not expected, a larger breeding season has been implicated in the evolution of mating strategies in *Phocidae* (*27*).

We suggest two avenues for future investigations specifically tailored to the hypotheses that follow from our results. One way to study mating behavior is through experimentally controlled environments, with known space and number of individuals, in a setup like the one used by a few recent studies with prairie voles (*10, 25*) and snapping shrimps (*26*). With this setup, an experimenter could vary the ratio of males and females, how long the animals are allowed to breed and the area available to test whether the predictions of our model hold empirically. This would be the live equivalent of our model, allowing the experimenter to see the magnitude of the effect of measured changes in the evolution of mating behavior. While a systematic investigation of several parameters would be unfeasible in this setup due to resource limitations, this would be an important step to check if our model’s predictions hold in a real environment. Another way to test the consequences of our model is using observational data from wild animals – which has been done before (*7*, *28*) – to see if a combination of OSR, season duration and female density explain the variation in mating behavior as it does in our simulation, irrespective of extrapair copulation, since systematic data on sexual fidelity among different species is harder to obtain.

Although the model was conceived with mammals in mind, there is nothing in it that makes it applicable exclusively to mammals. For example, our model may also explain the high prevalence of social monogamy in birds (*29*), since reported OSRs of bird populations are usually male-biased and females are more dispersed, which has been implicated as driving cooperative breeding (*31*), in a situation where our model consistently predicts monogamy.

## Methods

### Statistics

For the results in Table 2, percentage monogamy was regressed on female radius, season duration, number of males and a dummy variable for cheating (1 if it was present, 0 otherwise). All variables were normalized to be between 0 and 1 prior to regression. All statistical analysis and plotting was performed in R 3.4.0 (2). Raw data analyzed for the regression and figures is available as a Supplementary Data File (S1).

**Table 1.** Summary of regression results.

|  | **Estimate** | **Std. Error** | **t value** | **p-value** |
| --- | --- | --- | --- | --- |
| *Female Spacing* | 0.31006 | 0.01658 | 18.695,000 | <2e-16 |
| *Season Duration* | -0.28848 | 0.01483 | -19.447,000 | <2e-16 |
| *Number of Males* | 0.60222 | 0.01354 | 44.472,000 | <2e-16 |
| *Cheating* | -0.18516 | 0.01106 | -16.746,000 | <2e-16 |

Multiple R-squared = 0.7815; Adjusted R-squared = 0.7804

**Table 2.** Description of model parameters and simulated values.

| Parameter | Values | Description |
| --- | --- | --- |
| pregnancy-chance | 5 | The chance that a female will get pregnant in any given copulation. |
| number-of-males | 15; 20; 25 | The number of male agents in the simulation. The proportion between males is one of the main parameters we vary and they largely influence the outcome. |
| longevity | 4 | The number of breeding seasons a male agent survives - reflecting how many years males of a species are sexually active. We set it at 4 seasons, because increasing it makes the simulations much longer and more computationally costly. This parameter does not seem to qualitatively change the results, except in exacerbating whatever differences exist between the fitness of the two phenotypes. |
| number-of-females | 20 | The number of female agents in the simulation. |
| female-radius | 5; 10; 15; 20; 25; 30; 35 | This defines how much space there is between females. They are all disposed equally spaced in a rectangular grid. This is a parameter we specifically vary in every simulation, since this is of concern to the female dispersion hypothesis. |
| season-duration | 200; 400; 600; 800 | Defines how many ticks a breeding season lasts. This is grounded on the fact that males move one patch per tick. In our experiments, the total duration of the simulations was usually set at 150 generations, which makes up to 150 x longevity x season-duration ticks. This is usually set at 300 ticks, but varying it has a predictable and interesting effect on the outcome. |
| refractory-period-duration | 10; 30 | How long after copulation it takes for a female to be available again. This plays a dual role in the simulation. First, it limits the rate at which a monogamous male copulates - since his monogamous female partner is always available to him. Second, it prevents the polygamous male from behaving as a monogamous one, by not allowing him to copulate repeatedly with the same female - since there's a refractory period, he'll necessarily go after other available females during this time. |
| cheating-allowed | true; false | Whether monogamous males move around when their female partner is refractory; if true, it allows a large window for extrapair copulation. |
| percentage monogamy | outcome | The percentage of breeding season in which more than half the population of males was monogamous. |

### Software and Code Availability

The agent-based model was implemented using the software NetLogo (version 5.3.1) (1). The code is available as a Supplementary Data File (S2).

### Spatial distribution of agents

The whole field was set as a square grid with enough patches to fit all females, regularly spaced. Female agents remain fixed, regularly spaced according to the female radius parameter. Male agents are dispersed randomly between the female agents at the beginning of each breeding season.

### Male behavioral strategy

During each breeding season, a male moves one patch per tick – the measure of time within NetLogo. The behavior of the male agent depends on whether its genotype is monogamous or polygamous. Initially, both monogamous and polygamous males search around for a female, going after the one closest to their position that is *available*, i.e., a female which is neither pregnant nor refractory nor has a male closer to her. In case no female fulfills these criteria, the male moves to a random neighbor patch. If an available female is found, the male will go towards her. Upon occupying the same patch, the male copulates with the female, which then either gets pregnant – thus making her unavailable to other males until the end of the current breeding season – or enters the refractory period – a short period during which the female rejects all males (even her monogamous partner).

After copulation, the behavior of monogamous and polygamous males diverges. If cheating is not allowed, monogamous males will stay beside the first female with which they copulate, effectively making her unavailable to other males. However, that might limit his own chances of reproduction. Polygamous males, on the other hand, after copulating, return to their initial behavior, restarting the search for an available female.

We model males as having an underlying tendency towards monogamous or polygamous behavior. We assume this tendency to be genetic. This might not be realistic, as a change of environment and social conditions (such as being raised in captivity) might change mating behavior. However, recent studies are starting to bridge the gap between mating behavior, ecological features and its underlying neurological and genetic aspects (*25*), which suggests our assumption is not one that compromises the conclusions.

### Pregnancy and births

If a female agent gets pregnant, as mentioned before, it stays pregnant until the next breeding season. The simulation keeps track of a gene pool (called *trait pool* in the model). At the end of the breeding season, each pregnant female contributes to the gene pool, adding either a monogamous or a polygamous progeny to the gene pool. Note that these will not be in the simulation immediately, but only after the end of this generation, when all the current males die and are replaced with new ones – while artificial, this is done for pragmatic reasons, to keep the male population constant.

Whether a pregnant female will have a monogamous or polygamous progeny will depend on the genotype of the male agent who is the father. In the implementation, monogamy is represented by a value of 1 and polygamy by 0. The progeny’s genotype (G1) is determined as follows from the genotype of the father (G0):

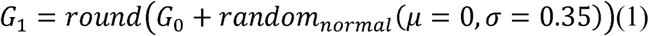

where ***round*** is a custom rounding function – any argument larger than or equal to 0.5 gives 1, otherwise it gives 0 – and ***random_normal*** is a function which returns a value randomly drawn from a normal distribution with mean and standard deviation specified by the arguments. This variation upon the father’s genotype is added to prevent the population from being stuck in a particular configuration if all agents have the same genotype. The first generation of males has an equal chance of being either monogamous or polygamous.

### Life cycle and breeding season

Each male agent survives for a number of reproductive seasons, after which they die. The population of males is kept constant, so when a male dies, a new one is added to the population at a random spot, whose mating behavior is drawn randomly from the gene pool. This means that not every pregnancy will lead to a new male with a given genotype, causing the distribution of monogamy/polygamy in the population to lag the underlying gene pool distribution. However, we see in our simulations that this does not change the results qualitatively – at the limit, for a high enough number of seasons, they are equivalent. The population of females is also kept constant. This can be thought of as assuming the number of males and females available for reproduction every breeding season does not vary a lot.

We explicitly disregard the non-breeding periods. We take the model to represent only the males and females within reproductive age and only during the times where females are receptive to mating attempts by the males.

## Author Contributions

All authors contributed equally to this work. K.N. implemented the model. B.V.G., D.M.G., D.S. and K.N. designed the model and experiments, analyzed the results and wrote the paper.

## Acknowledgements

We would like to acknowledge the funding agency Conselho Nacional de Desenvolvimento Científico e Tecnológico (CNPq). We thank Jonas, Júlia and Pamela for all support given. We also thank Dieter Lukas for all the discussion and suggestions. The authors declare no conflict of interest.

## References

1. S. R. Tecot, B. Singletary, E. Eadie, Why “monogamy” isn't good enough: Pair-Living, Pair-Bonding, and Monogamy. Am. J. Primatol. 78, 340–354 (2016).

2. P. J. Greenwood, Mating systems, philopatry and dispersal in birds and mammals. Anim. Behav. 28, 1140&1162 (1980).

3. D. G. Kleiman, Monogamy in mammals. Q. Rev. Biol. 52, 39&69 (1977).

4. C. Borries, T. Savini, A. Koenig, Social monogamy and the threat of infanticide in larger mammals. Behav. Ecol. Sociobiol. 65, 685–693 (2011).

5. C. Opie, Q. D. Atkinson, R. I. M. Dunbar, S. Shultz, Male infanticide leads to social monogamy in primates. Proc. Natl. Acad. Sci. 110, 13328–13332 (2013).

6. D. Lukas, T. H. Clutton-Brock, The Evolution of Social Monogamy in Mammals. Science. 341, 526–530 (2013).

7. P. E. Komers, P. N. Brotherton, Female space use is the best predictor of monogamy in mammals. Proc. Biol. Sci. 264, 1261–1270 (1997).

8. R. Hilgartner, C. Fichtel, P. M. Kappeler, D. Zinner, Determinants of Pair-Living in Red-Tailed Sportive Lemurs (Lepilemur ruficaudatus): Pair-Living in Red-Tailed Sportive Lemurs. Ethology. 118, 466–479 (2012).

9. L. M. Porter, Social organization, reproduction and rearing strategies of Callimico goeldii: new clues from the wild. Folia Primatol. 72, 69–79 (2001).

10. A. G. Ophir, S. M. Phelps, A. B. Sorin, J. O. Wolff, Social but not genetic monogamy is associated with greater breeding success in prairie voles. Anim. Behav. 75, 1143–1154 (2008).

11. K. E. Mabry, E. L. Shelley, K. E. Davis, D. T. Blumstein, D. H. Van Vuren, Social Mating System and Sex-Biased Dispersal in Mammals and Birds: A Phylogenetic Analysis. PLoS ONE. 8, e57980 (2013).

12. D. Ren et al., Genetic Diversity in Oxytocin Ligands and Receptors in New World Monkeys. PLoS ONE. 10, e0125775 (2015).

13. L. Dieter, T. Clutton-Brock, Evolution of social monogamy in primates is not consistently associated with male infanticide. Proc. Natl. Acad. Sci. 117, e1674 (2014).

14. C. Opie, Q. D. Atkinson, R. I. Dunbar, S. Shultz, Reply to Lukas and Clutton-Brock: Infanticide still drives primate monogamy. Proc. Natl. Acad. Sci. 111, e1675–e1675 (2014).

15. S. E. Ptak, M. Lachmann, On the evolution of polygyny: a theoretical examination of the polygyny threshold model. Behav. Ecol. 14, 201–211 (2003).

16. S. Gavrilets, Human origins and the transition from promiscuity to pair-bonding. Proc. Natl. Acad. Sci. 109, 9923–9928 (2012).

17. R. Schacht, A. V. Bell, The evolution of monogamy in response to partner scarcity. Sci. Rep. 6, 32472 (2016).

18. S. M. Henson, J. L. Hayward, The mathematics of animal behavior: An interdisciplinary dialogue. Notices Amer. Math. Soc. 57, no. 10 (2010).

19. H. V. D. Parunak, R. Savit, R. L. Riolo, Agent-based modeling vs. equation-based modeling: A case study and users' guide. MAS&S. 10–25 (1998).

20. D. L. DeAngelis, W. M. Mooij, Individual-based modeling of ecological and evolutionary processes. Annu. Rev. Ecol. Evol. Syst. 36, 147-168 (2005).

21. C. T. Bauch, R. McElreath, Disease dynamics and costly punishment can foster socially imposed monogamy. Nat. Commun. 7, 11219 (2016).

22. K. Annaliese et al., An evolutionary framework for studying mechanisms of social behavior. Trends Ecol. Evol. 29, 581&589 (2014).

23. K. Sharma et al.., Vigorous dynamics underlie a stable population of the endangered snow leopard *Panthera uncia*in Tost Mountains, South Gobi, Mongolia. PLoS ONE. 9, e101319 (2014).

24. I. Herfindal et al.., Population properties affect inbreeding avoidance in moose. Biol. Lett. 10, 20140786 (2014).

25. M. Okhovat et al., Sexual fidelity trade-offs promote regulatory variation in the prairie vole brain. Science. 350, 1371-1374 (2015).

26. L. M. Mathews, Tests of the mate-guarding hypothesis for social monogamy: male snapping shrimp prefer to associate with high-value females. Behav. Ecol. 14, 63&67.

27. J. F. Wtittenberger, R. L. Tilson, The evolution of monogamy: Hypotheses and evidences. Annu. Rev. Ecol. Syst. 11, 197&232 (1980).

28. F. S. Dobson, B. M. Way, C. Baudoin, Spatial dynamics and the evolution of social monogamy in mammals. Behav. Ecol. 21 (04), 747&752 (2010).

29. N. L. Ford, Variation in mate fidelity in monogamous birds. in Current Ornithology Ed. Richard F. Johnston (Springer, US, 1983). 329&356.

30. P. F. Donald, Adult sex ratios in wild bird populations. Ibis. 149, 671&692 (2007).

31. J. Komdeur, T. Székely, X. Long, S. A. Kingma, Adult sex ratios and their implications for cooperative breeding in birds. Philos. Trans. R. Soc. Lond. B. Biol. Sci. 372, 20160322 (2017).

## Methods -References

32. U. Wilensky. NetLogo. http://ccl.northwestern.edu/netlogo/ Center for Connected Learning and Computer-Based Modeling, Northwestern University (1999).

33. R Core Team. R: A language and environment for statistical computing. https://www.rproject.org/. R Foundation for Statistical Computing, Vienna, Austria (2017).

